# Genome of the enigmatic watering-pot shell reveals adaptations of a living suction anchor

**DOI:** 10.1101/2024.10.08.617303

**Authors:** Julia D Sigwart, Nur Leena W.S. Wong, Vanessa Liz González, Fabrizio Marcondes Machado, Carola Greve, Tilman Schell, Zeyuan Chen

## Abstract

The watering pot shells have been rightly called “the weirdest bivalves” for their highly modified body plan and fused tubular shell that resemble the spout of a watering can. This adventitious tube has defied rational explanation as an adaptive pathway and yet arose twice convergently. We present the first genome for the clade Anomalodesmata: an annotated chromosome-scale genome of *Verpa penis* (Linnaeus, 1758). The first watering pot shell ever described ranges across southeast Asia but is nonetheless extremely rare and has never previously been sequenced. The assembly length of 507 Mb, with a contig N50 of 5.33 Mb, has 96.5% of sequences anchored onto 19 pseudochromosomes. Contrary to expectations from such a highly modified body plan, there is no evidence of chromosome reduction compared to the ancestral condition of heterodont bivalves (1N=19). A new hypothesis based on analysis of the living animals and literature explains the adaptive significance of this body form: it is structurally optimised for vertical stability in soft sediments, with parallels to engineering principles of a suction anchor. Our study offers new insights to a long-standing mystery in molluscan body forms, and provides genomic resources that are relevant to understanding molluscan evolution, biomineralisation, and biomimetic design.

## Introduction

Bivalvia, a diverse and widespread molluscan clade, is widely recognised for environmental and economic significance. Locally abundant species of mussels and oysters are frequently credited for “ecosystem services” such as water filtration and as indicators of ecosystem health[1]. The deep fossil record of bivalves is a key data source for understanding the impact and recovery around past mass extinction events[2]. But bivalves are neither ecologically nor morphologically homogenous, and the clade contains frequent and repeated body plan modifications[3], such as vermiform ship worms and the mysterious watering-pot shells.

The superfamily Clavagelloidea (Anomalodesmata) includes two families whose species produce a tubular adventitious shell. It widens at the base and the bottom is enclosed with a plate perforated by holes that resemble the spout of a watering can (Fig 1). The animals of this form are immobile, buried in soft sediments, with the narrow, open end of the tube emerging slightly from the surface. The only resemblance to a bivalve shell is the larval shell retained on one (dorsal) side of the adult tube.

**Figure 1.**
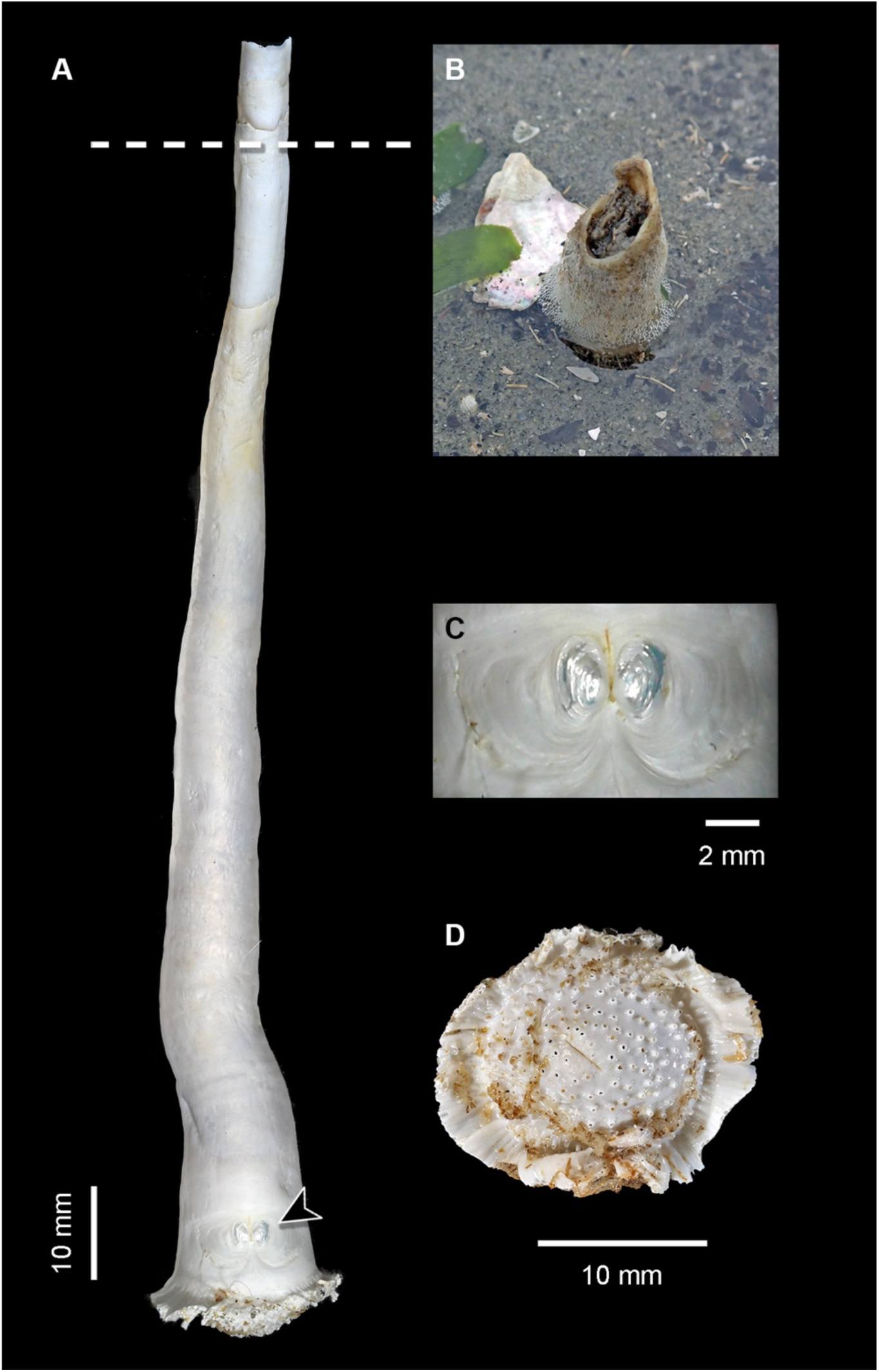
*Verpa penis*, the watering pot shell, from Merambong Shoal, Johor, Malaysia. A, animal in dorsal view, with dashed line indicating typical burial depth, B, *in situ* image of live animal, photo by NLWSW, C, close up of the larval shell (indicated by an arrowhead in A), D, anterior (bottom) “watering pot”. Photos (except B) of specimen SMF 367281 by Sigrid Hof.

The two families, Clavagellidae and Penicillidae, are morphologically and anatomically distinct[4]. Clavagellidae retain only the left valve within the fused tube or crypt, whereas Penicillidae retain both valves; this correlates to differences in the musculature. Both families include species with endobenthic, tube-forming bodies, but also other epibenthic crypt-forming genera[4,5], meaning that this form has evolved twice independently[6].

Clavagelloidean anatomy and shells show significant modifications compared to the canonical bivalve form. The body is positioned at the bottom end of the tube (anatomically anterior), with a pedal disc in contact with the perforated “watering pot” and siphons extending upward through the vertical tube. The tube forms as post-settlement growth of the larval shell, which expands in the normal way but at some point fuses[4]. The term ‘adventitious’ refers to the unusual anatomical position rather than accidental or externally moderated formation. The adventitious tube is traditionally described as a defensive adaptation[5,7]; it provides armour protecting the siphons, but this hypothesis has never been scrutinised. The presence of arenophilic (detritus-sticking) glands in the siphon tips of some clavagelloids[7] weakens this defensive idea, as the glands provide camouflage that appears to have the same anti-predation function.

Formal morphocladistic analyses support the convergent evolution of this remarkable structural change[8]; however, molecular data are scarce for either family. To date, sequence data are only available for three genera in both families: coral-boring *Bryopa* (Clavagellidae), including one mitogenome[9–11], and some standard markers for adventitious tube-forming *Brechites* and *Verpa* (as *Penicillus*) (Penicillidae)[9,12]. Further data for any watering pot shell would allow insights to the potential genomic mechanisms underlying this extreme adaptation.

## Methods

### Field collection

Animals were collected at Merambong Shoal, Johor, Malaysia, 11 February 2020 (1.3394°N, 103.6066°E) and maintained alive in laboratory culture at I-AQUAS, University Putra Malaysia, Port Dickson, for several months. One female individual was dissected into separate organs, with the foot and siphons preserved in ethanol and other organs preserved in RNAlater, this individual was used for all aspects of genome assembly (PacBio, Hi-C, RNAseq). The remaining specimens from the same collecting event are retained in the University Putra Malaysia Marine Collection (UPMMC), Port Dickson.

### Sequencing

High molecular weight (HMW) gDNA was extracted from ethanol-preserved tissues[13] with a pre-wash step with sorbitol. DNA concentration and DNA fragment length were assessed using the Qubit dsDNA BR Assay kit on the Qubit Fluorometer (Thermo Fisher Scientific) and the Genomic DNA Screen Tape on the Agilent 2200 (Agilent Technologies). We prepared one low-input PacBio HiFi library according to the SMRTbell Express Prep Kit v2.0 instructions. To remove smaller fragments, the DNA library was size selected using beads with a cut-off at 3kb.. The same library was loaded on two SMRT 8M cells and sequenced in CCS mode using the PacBio Sequel II instrument. The on-plate concentration was 80 pM using adaptive loading and the Sequel II Binding kit 2.2 (Pacific Biosciences, Menlo Park, CA). Pre-extension time was 2 hours, run time was 30 hours. HiFi reads were called using a pipeline running PacBio tools: ccs 6.4.0, actc 0.3.1, samtools 1.15[14], and DeepConsensus 1.2.0[15].

To prepare a chromatin conformation capture library, we used the Arima High Coverage Hi-C Kit v01 (Arima Genomics) according to the Animal Tissue User Guide for proximity ligation using approximately 44 mg of muscle tissue. The proximally-ligated DNA was then converted into an Arima High Coverage HiC library according to the Swift Biosciences Accel-NGS 2S Plus DNA Library Kit protocol. The fragment size distribution and concentration of the Arima High Coverage HiC library were assessed using the TapeStation 2200 (Agilent Technologies) and the Qubit Fluorometer and Qubit dsDNA HS reagents Assay kit (Thermo Fisher Scientific, Waltham, MA), respectively. The library was sequenced on the NovaSeq 6000 platform at Novogene (UK) using a 150 paired-end sequencing strategy, resulting in an output of 21 Gb.

Total RNA was isolated from pedal disc, mantle, siphon and pericardium tissues using TRI-Reagenz® (Sigma-Aldrich) according to the manufacturer’s instructions. The quality and concentration of each extraction were assessed using the TapeStation 2200 (Agilent Technologies) and the Qubit Fluorometer with the RNA BR Reagents Assay Kit (Thermo Fisher Scientific, Waltham, MA). The RNA extractions were then pooled at equal concentrations and sent to Novogene (UK) for Illumina paired-end 150 bp RNA-seq of a cDNA library (insert size: 350 bp) with an expected output of 10 Gb.

Genome size and heterozygosity were estimated from a k-mer profile of the HiFi reads (Fig S1). First, a count from Jellyfish 2.3.0[16] was run with the additional parameters “-m 21 -s 100M” and all HiFi reads as input, then the GenomeScope v1.0 and v2.0[17] in combination with R 4.3.1 was executed with the k-mer size set to 21.

All the HiFi reads were processed for *de novo* genome assembly using hifiasm v0.14-r312 with default parameters[18]. To remove haplotypic duplications and overlaps in the assembly, Purge_Dups v.1.2.5 was used[19]. The Hi-C reads (145,310,480 reads, length 150 bp, total data size of 21,796,572,000 bp) were then aligned to the initial genome assembly using Arima Genomics’ mapping pipeline (https://github.com/ArimaGenomics/mapping_pipeline). In short, the reads were mapped to the reference using BWA-MEM v0.7.17-r1188[20], converted to a sorted .bam file, and filtered to keep uniquely mapping pairs. PCR duplicates were removed using Picard v3.0.0[21]. The final alignment file and the assembly were then passed to YaHS v 1.1a-r3[22] for scaffolding in the default mode. Juicebox Assembly Tools[23] were used to generate and visualise a HiC contact map. We manually curated the scaffolded assembly using an editable Hi-C heatmap to improve the assembly quality and to correct misassembles with Juicebox v1.11.08[24] (Fig S2).

Genome completeness was assessed by BUSCO v5.4.3, in euk_genome_met mode[25] using the lineage dataset metazoa_odb10 (Creation date: 2021-02-17, number of genomes: 65, number of BUSCOs: 954). (Table S1-S2). Completeness regarding k-mers and QV values were obtained with Meryl 1.3 and Merqury 1.3[26].

### Genome annotation

TRF v4.09[27] was used for tandem repeats identification. Transposable elements (TEs) were annotated using a combination of *ab initio* and homology-based approaches. First, repeat elements were identified *de novo* using RepeatModeler v2.0.4[28]. The predicted models, together with a repeat database Dfam_3.0, were then merged together and used as a custom library for RepeatMasker v4.1.5[28] to localise, identify and mask repeats.

Protein-coding genes were predicted using the following approaches: *ab initio* prediction, homology-based prediction, and transcriptome-based prediction. Around 79M RNA-seq reads were aligned to the *V. penis* genome using STAR v2.7.3a[29]. The resulting alignment file served as an important support in all three prediction methods. *Ab initio* gene prediction was performed on the soft repeat-masked assembly with Braker v3.0.3[30] using default parameters. Two closely related species, *Dreissena rostriformis*[31] and *Sinonovacula constricta*[32] were selected and used for homology-based prediction. First, the protein sequences of *D. rostriformis* and *S. constricta* were downloaded from NCBI and aligned against the assembled genome using MMseqs2[33] with the parameter “-e 100.0 -s 8.5 -- comp-bias-corr 0 --max-seqs 500 --mask 0 --orf-start-mode 1”. The results of homologous alignments were then combined into gene models with the splice site identified using mapped RNA-seq data with GeMoMa v1.9[34] applying default parameters. Finally, the gene predictions were further sorted and filtered separately using the GeMoMa module GAF with default parameters. For the transcriptome-based prediction, the transcriptome of *V. penis* was assembled by both *de novo* and genome-guided approaches using Trinity v2.15.0[35]. The results were merged and passed to Program to Assemble Spliced Alignments (PASA) v2.5.2[36] for gene predictions. In the end, all the predictions were combined into consensus coding sequence models using EVidenceModeler v1.1.1[36], with the weighting of each method as “*ab initio* 1; homology-based 2, transcriptome-based 8”. In the end, gFACs v1.1.2[37] was used to remove incomplete gene models. The completeness of the gene set was accessed using BUSCO v5.4.3[19], protein mode using the lineage dataset metazoa_odb10. The predicted genes were functionally annotated by aligning them to the eggNOG (emapper v1.0.3) databases (2023_06_26) using diamond v0.9.30.131.

### Orthology Inference and Phylogenetic Analysis

Predicted peptide sequences from 30 additional molluscan genomes were used downstream phylogenetic analyses (Supplementary Table S5), representing 23 Bivalvia genomes plus one Cephalopoda, two Scaphopoda and four Gastropoda protein sets used as outgroups.

Orthology prediction was conducted in OMA v.2.4.1[38]. Orthologous groups (OGs) were aligned using MAFFT v.7.407[39] with the L-INS-i algorithm[40] and the command option “- leavegappyregion.” OG alignments were trimmed with GBLOCKS v. 0.91b[41] to cull regions of dubious alignment. An ortholog matrix was built using a custom python script (selectslice.py)[42] to include loci with a minimum of 80% (3379 genes; 848,293 amino acids) taxa occupancy and was concatenated using Phyutility[43]. Phylogenetic reconstruction was conducted in IQ-Tree v. 2.1.2[44] using ModelFinder[45] for model selection and ultrafast bootstrap[46] with 1000 replicates. Model finder best fit model based on Akaike Information Criterion (AIC), Corrected Akaike Information Criterion, and Bayesian Information Criterion (BIC), LG+F+R7, was used for phylogenetic tree reconstruction in IQ-Tree. Nodal support values are represented by maximum likelihood bootstrap proportions. Phylogenetic reconstruction and genome assembly were conducted on “Hydra”, the Smithsonian Institution High Performance Cluster (SI/HPC; doi.org/10.25572/SIHPC).

### Systematics

The taxonomy of *Verpa penis* has raised some specific challenges that have been raised repeatedly[4,11] and deserve a short note: this species was originally named *Sabella penis*, as Linnaeus interpreted the animal as a tube worm. The genus name was later revised to the genus *Penicillus* Bruguière, 1789 which is taxonomically invalid and a junior homonym of the same name used for polychaetes, an obsolete pre-Linnean name[47]. There is no conflict with a potential alternative use because the polychaete name is not in use; the family name Penicillidae is still valid and correct.

## Results & Discussion

Anomalodesmatans represent 22 living families, characterised by a wide array of highly specialised morphologies[9]. Among these highly adapted bivalves are not only the tube-dwelling clavagelloids but also a broad group of carnivorous species that are highly diversified in the deep sea[8,48]. Anomalodesmatans in general are distinctive, but rare, and often in inaccessible habitats. Prior studies have repeatedly noted that they are under-represented in global phylogenetic research[10]. This study represents the first genome for any member of this clade and an important resource for future comparative studies.

The final assembly for the genome of *Verpa penis* had a total length of 507 Mb, a contig N50 of 5.33 Mb, a scaffold N50 of 27.6 Mb, and with 96.5% of the sequences anchored onto 19 scaffolds (Fig 2, Supplementary Table S2). The whole-genome sequencing yielded in total 17Gb of HiFi data (Supplementary Fig S2), which corresponds to a theoretical coverage of 36x. A total of 928 (97.2%) BUSCO benchmark set were complete (Supplementary Table S1) and QV (54.7) and k-mer completeness (91.8%) also suggest a high-quality assembly (Supplementary Fig S3). Contiguity statistics are more than seven-fold higher compared to other bivalves (Supplementary Table S2). Approximately 45.26% of the genome was composed of repetitive elements (38.23% transposons; Fig 2, Supplementary Table S2). Gene annotation combining evidence from transcripts, homologous proteins, and *ab initio* prediction resulted in 25,135 predicted protein-coding genes with an average length of 9,113 bp; the average transcript length, exon number, average exon length, and intron length were comparable to the closely related species (Supplementary Tables S3-S4). Among the predicted protein-coding genes, 76.3% could be annotated; the gene set shows high completeness with 97.9% of complete BUSCOs (Supplementary Table S5).

**Figure 2.**
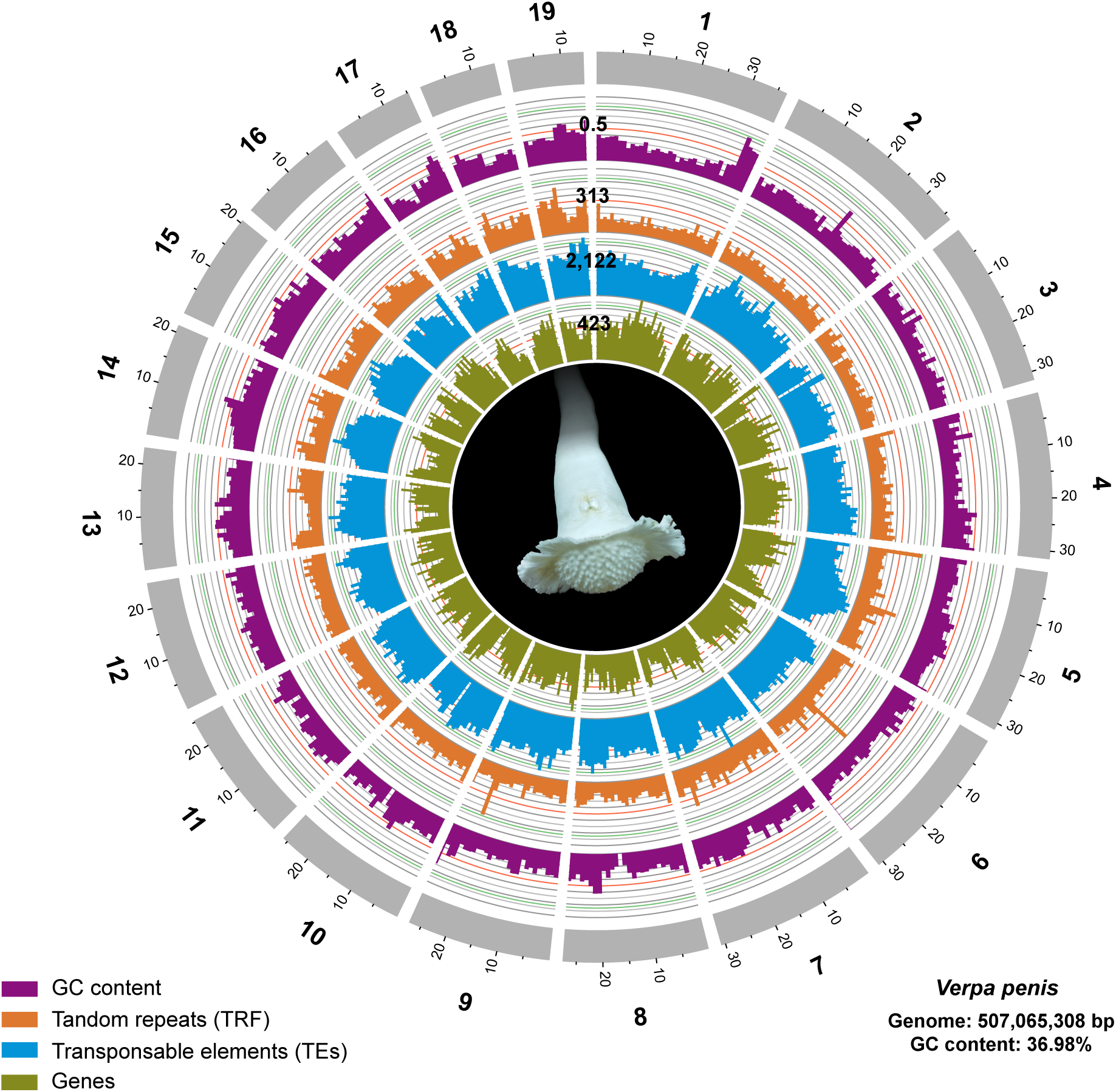
General characteristics of the *V. penis* genome. Tracks from inside to outside correspond to gene number, TEs number, TRF number and GC content in sliding windows of 1 Mb across each of the 19 pseudochromosomes, visualised by Circos v0.69-9. Photo by NLWSW.

The phylogeny confirms Anomalodesmata, represented by *Verpa*, is sister to Imparidentia (Fig 3); these two orders together form the clade Euheterodonta[49]. The earliest molecular analyses found Anomalodesmata inside the historical “Heterodonta”[9,12,49] although there were several exceptions with alternative topologies. Early morphological studies were unable to recover monophyletic Euheterodonta and several molecular studies proposed alternative placements for Anomalodesmata[50]. Nonetheless, studies that focussed on Anomalodesmata, and later larger scale multi-loci and transcriptome analyses refined the phylogenetic hypotheses to identify the families that comprise Euheterodonta[11]. The consistency of this result in several independent studies led to the recognition of the clade Imparidentia as the euheterodont bivalves excluding anomalodesmatans[51].

**Figure 3.**
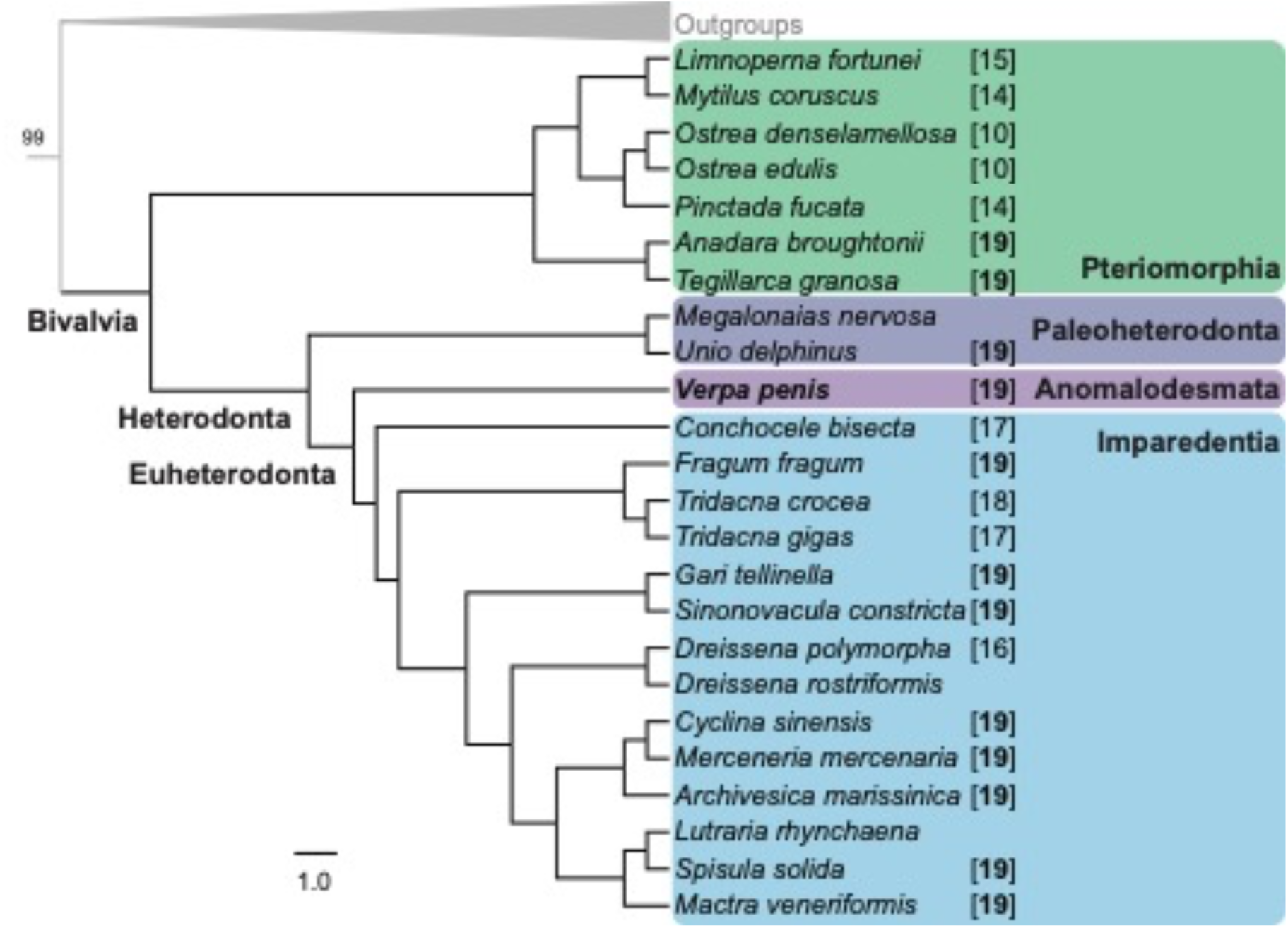
Phylogenetic placement of *Verpa penis*. Maximum likelihood topology generated in IQ-Tree using 80% minimum taxa occupancy matrix (3379 genes; 848,293 amino acids). Numbers in square brackets after the taxon names indicate 1N chromosome numbers.

Genome resources for molluscs remain scarce, for multiple reasons[52]. The diversity across bivalves is relatively better sampled than other molluscan clades, but there is a disproportionate focus on species of economic interest, which mainly fit in the clade Pteriomorphia including oysters, scallops, and mussels. By contrast, the heterodont bivalves, Euheterodonta, Archiheterodonta, and Palaeoheterodonta, comprise a larger fraction of species richness and broader variety of habitats including freshwater radiations, but lag behind in available genomes. We selected a subset of species for our analyses to represent a balance of diverse species and some adaptation to robustly test the placement of Anomalodesmata among other bivalve clades.

The ancestral genome for Imparidentia had a karyotype of 1N=19 [53], which is identical to other closely related clades, as known from Paleoheterodonta[54], and now one Anomalodesmata (Fig 3, Supplementary Table S5). Most likely this is the ancestral karyotype for all heterodont bivalves. Within Imparidentia, several lineages show chromosome reductions: in the freshwater invasive zebra mussels, and symbiont-hosting deep-sea *Conchocele*[55] and tropical photosymbiotic *Tridacna*[56,57]. *Verpa penis* is likewise highly adapted, and very rare, but has a unreduced karyotype.

Genome re-arrangements such as chromosomal inversions and chromosome fusion are invoked in explaining adaptive processes[58,59]. Genome reductions have consistent patterns in the evolution of mammals[60] but this is evidentially not universal. The retention of a full set of chromosomes in *Verpa* does not refute the general idea that species with smaller populations or those in isolated environments might benefit from chromosomal reductions that preserve genetic stability and reduce variation; however, it indicates these are not universal mechanisms.

Many other bivalves have burrowing or boring adaptations, usually in harder substrata, which have emerged convergently at least eight times, but most of them look like typical bivalves; the adaptations of Clavagelloidea are so extreme they cannot be included in morphometric analyses of shell shape[3]. The adventitious tube is not truly endolithic but grows in a matrix of relatively soft substrata, in sand or mud. For this reason, clavagellids can be classified either tube dwellers in soft sediments or facultative semiendolithic borers[61]. Observations of living animals are rare, but it is apparent that specimens removed from the substrate generally cannot rebury themselves, and if they stay safe in a prostrate position may even start growing a tube bent upwards[62]. As long as the anterior watering pot is submerged in the sediment, however, they can likely manoeuvre themselves back into normal vertical position.

The perforations of the bottom watering-pot disc can be closed by the pedal disc, or opened to allow the flow of water through the tube. Mixing is important to avoid stratification of an oxygen gradient within the enclosed tube, and an essential component of the limited mobility for these animals. Observations of *Verpa penis* gave some insight to the pedal disc contractions that pump interstitial water out of the mantle cavity: the anterior part of the mantle and the pedal aperture operate like the piston and valve of a pump[63]. By generating contractions that pump water across the basal plate, the bivalve is apparently able to liquify the surrounding sediment, reducing its density and resistance[4]. This liquefaction not only allows the bivalve to sink into the substrate but would also aid in maintaining its position within the sediment.

Molluscan evolution encompasses multiple strategies for animals to maintain position and defend themselves from disturbance. Bivalves might increase size and mass, to achieve deeper burial and avoid disturbance[64]. Other adaptations include engineering modifications of the sculpture to create a ratchet structure[65], or weight distribution within shells to passively maintain position[64,66]. While *Verpa penis* has a relatively smooth shell, other tube-forming clavagelloideans have ornamentation (e.g. siphonal collars, anterior tubules)[4] that likely functions as stabilisers and to increase surface area. These may be related to the specific sediment properties of the habitat for each species. For example, *Verpa* has an anterior fringe, missing in some other genera, that extends the surface area of the basal plate and has been interpreted as an adaptation for burrowing[64].

The watering pot shell appears to follow engineering principles that optimise suction resistance, similar to those used in suction caisson anchors. Mud creates a suction force that can pull objects down, making them hard to lift or remove, especially if there is a large vertical surface area. However, a long, thin object submerged in mud is not necessarily stable against tipping. An enlarged, cone-shaped or curved base stabilises the structure. These principles mean that suction caissons, or adventitious tubes, once in place, are stable and resistant to lateral forces (such as currents or winds) because of the large surface area in contact with the mud. The overall form is stable and also resistant to removal, which is a significant advantage for a sessile bivalve, exploiting the natural suction property of soft sediment. But when vertical movement is needed, this can be enabled by a valve at the bottom of a suction caisson or adventitious tube: moving fluid via air pockets or perforations, such as the basal plate, breaks the suction seal and allows up or down movement as needed. Anatomical modifications observed in most adult clavagelloids—such as a reduced foot, degenerated adductor muscles, and especially the absence of byssus threads[67]— reinforce the plausibility of the anchor-tube theory. Over time, the adventitious tube has taken on another major role of the shell: acting as an exoskeletal attachment site for the reduced musculature[7]. The whole watering pot structure—including the adventitious tube as an intrinsic extension—enables efficient burrowing, protection, and stabilisation in soft marine sediments.

This study benefitted from an unusually high number of specimens found in a single collecting event, where we were able to find six individuals. Conditions during that event were abnormal, with heavy rain producing a freshwater layer on top of a tidally exposed shoal, and the animals were found by tactile searching rather than visually. This opportunistic collection may reflect circumstance where the sediment was liquified and shifting around the animals such that they could not reposition themselves vertically.

It is clear that this species is rare, as it typical of all Clavagelloidea[4], and highly vulnerable to environmental disturbance that would shift sediment profiles. The population in Johor, Malaysia, is potentially threatened by anthropogenic activities through active land reclamation, and it is also geographically adjacent to a population in Singapore that was recently rediscovered after being reported as locally extinct[68].

In evolutionary biology we frequently seek to explain adaptations, or at least classify them in terms of determinism (inevitable consequence of functional need) or contingency (many different ways to solve the same problem). The convergence of adventitious tubes in clavagelloidean bivalves is apparently a deterministic outcome of optimising stability in soft sediments, and parallels engineering principles of suction caissons. The genome of *Verpa penis* demonstrates that such remarkable modification in body plan does not correlate to significant changes in genome architecture. These new results provide a new basis for further understanding of the “biomineralistion toolkit” and the fascinating ecology of these vulnerable animals.

## Acknowledgements

This project was funded by the LOEWE Translational Biodiversity Genomics (TBG) and by the Leibniz Association project PHENOME. We are grateful to many colleagues for their support for this work including technical assistance, field work, and fruitful discussion, in particular Ekin Tilic for taxonomic advice. We thank Damian Baranski, Alexander Ben Hamadou, and Charlotte Gerheim, for support with lab work and sequencing. The Genome Technology Center (RGTC) at Radboudumc and Sequencing Core Facility (Nijmegen, The Netherlands) provided the PacBio SMRT sequencing service on the Sequel II platform. This is contribution number 43 of the Senckenberg Ocean Species Alliance.

## Data and materials availability

The whole genome sequencing data and novel genome assembly of *Verpa penis* are available via the National Center for Biotechnology Information (NCBI) under the PRJ number PRJNA1120794.

## Supplementary results

Additional genome measures (Figures S1-S3, Tables S1-S4) are summarised below for *Verpa penis*, as well as the source data for the additional species used in the phylogenetic reconstruction (Table S5).

**Figure S1.**
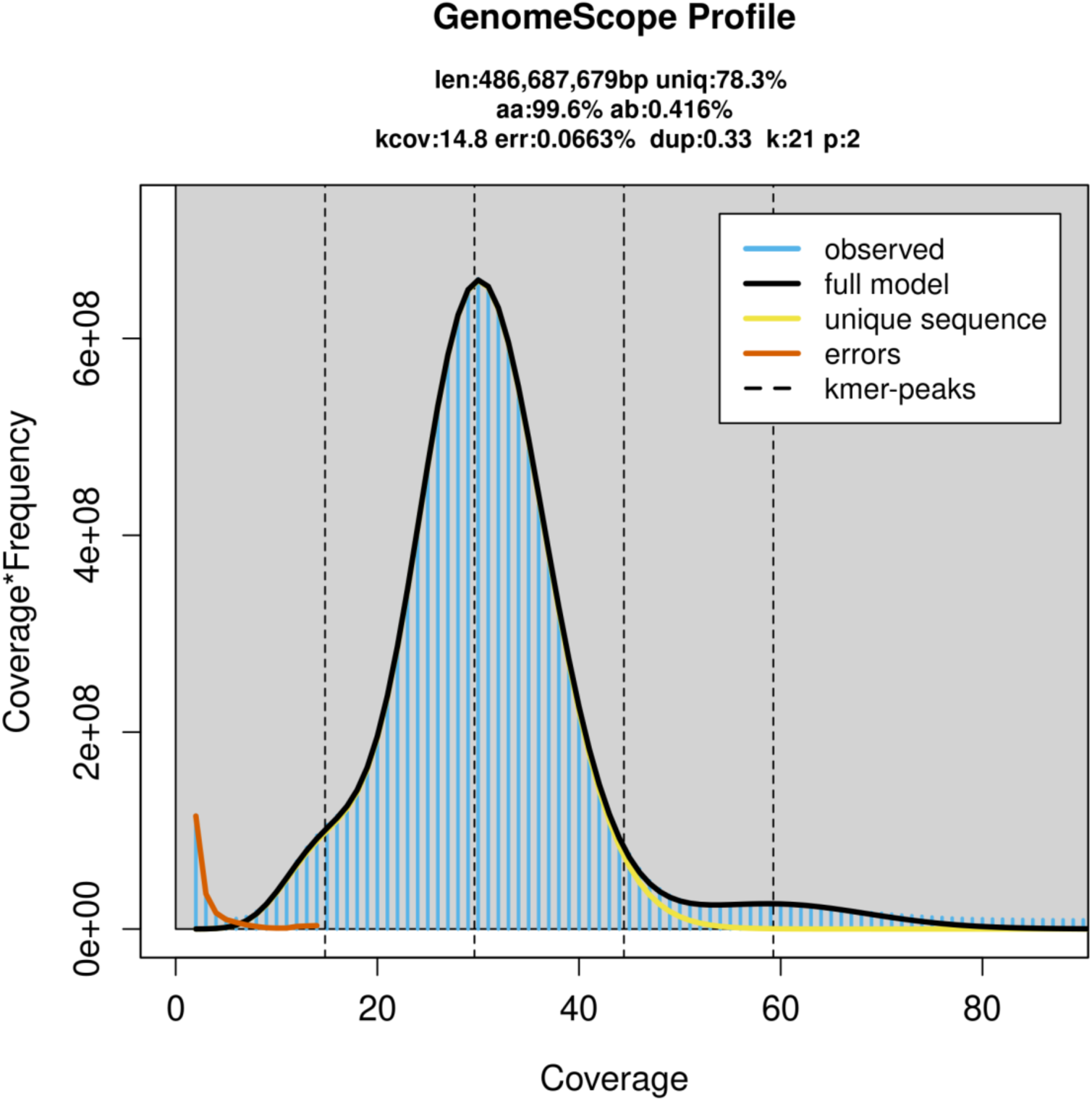
Estimation of genome features based on the distribution of 21-mer frequency.

**Figure S2.**
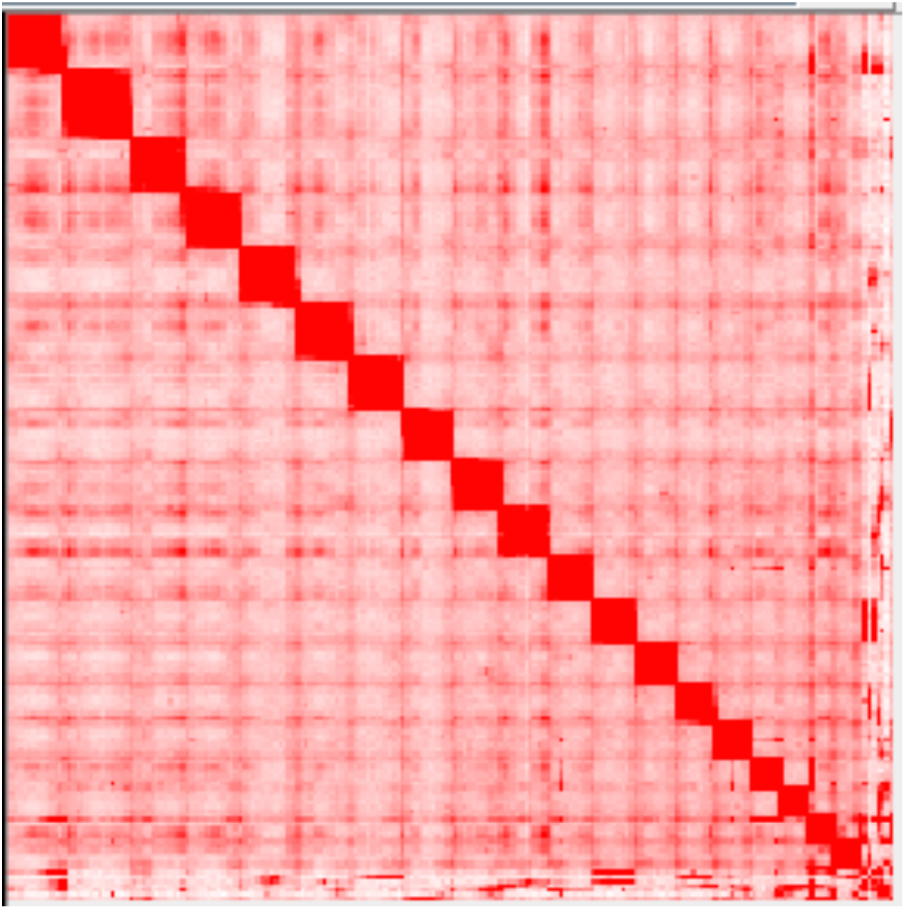
Hi-C chromosome contact maps. Each block represents a Hi-C contact between two genomic loci within a 1 Mb window. The darker the shade of a block, the higher the contact intensity.

**Figure S3.**
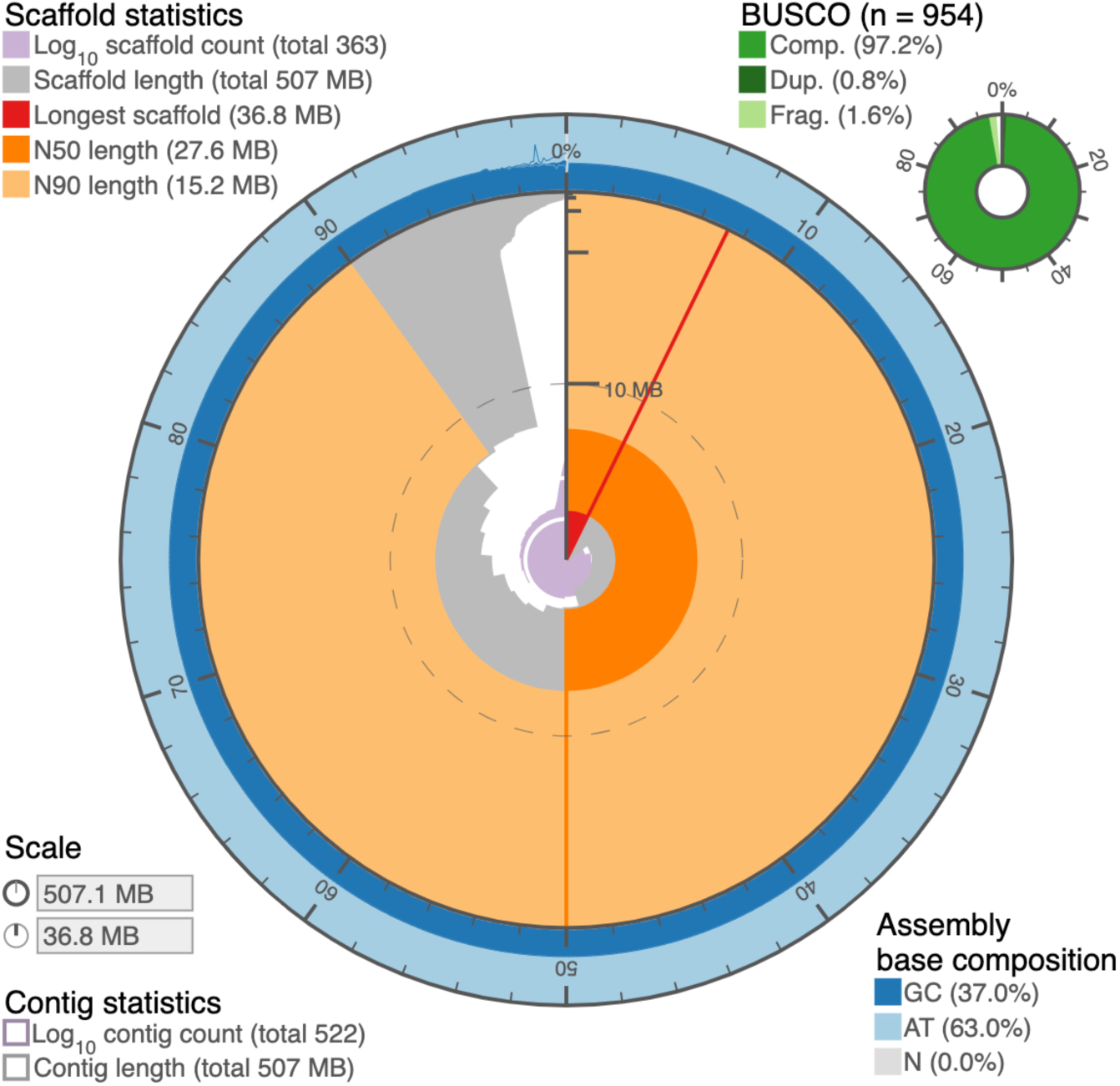
Snail plot summarizing the genome statistics of *Verpa penis*. Snail plots were generated using BlobToolKit v 3.3.10 (Challis et al., 2020). The distribution of scaffold lengths is shown in dark gray with the plot radius scaled to the longest sequence present in the assembly (36.8Mb, shown in red). Statistics of note shown here include the genome N50: 27.6 Mb (dark orange), N90: 15.2Mb (light orange),and base composition (percentage of GC in dark blue, AT in light blue, and N in light gray). BUSCO results summarizing complete, duplicated and fragmented using the Mollusca_odb10 dataset are displayed in top right (in shades of green).

**Table S1.** Statistics of the final assembly at different assemble stages.

| Characteristic | Value |
| --- | --- |
| Total Contig length (bp) | 507,038,511 (522) |
| Contig N50 (bp) | 5,325,940 (28) |
| Longest Contig (bp) | 16,098,946 |
| Total scaffold length (bp) | 507066948 (636) |
| Scaffold N50(bp) | 27,570,495 (8) |
| Longest Scaffold (bp) | 36,765,564 |
| N's per 100 kbp | 5.61 |
| Complete BUSCOs | 97.2% (928) |
| Complete and single-copy BUSCOs | 96.4% (920) |
| Complete and duplicated BUSCOs | 0.8% (8) |
| Fragmented BUSCOs (F) | 1.6% (15) |
| Missing BUSCOs (M) | 1.2% (11) |

**Table S2.** Identified repeat classes in the *V. penis* genome.

| Type | Repeat Size(bp) | % of Genome |
| --- | --- | --- |
| Tandem repeats | 35,671,495 | 7.03 |
| Transposable elements (total) | 193,845,983 | 38.23 |
| DNA | 46,038,611 | 1.03 |
| LINE | 13,410,503 | 2.64 |
| LTR | 7,525,767 | 1.48 |
| Low | 875,480 | 0.17 |
| PLE | 643,942 | 0.13 |
| RC | 17,276,857 | 3.41 |
| Retroposon | 1,216 | 0.00 |
| SINE | 25,028,422 | 4.94 |
| Satellite | 126,310 | 0.02 |
| Unknown | 95,656,288 | 18.86 |
| scRNA | 835 | 0.00 |
| srpRNA | 3,291 | 0.00 |

**Table S3.** Statistics of predicted protein-coding genes in the *V. penis* genome.

| Gene set | Total Genes Predicted | Average Transcript Length(bp) | Average CDS Length(bp) | Average Exon per Gene | Average Exon Length(bp) | Average Intron Length(bp) |
| --- | --- | --- | --- | --- | --- | --- |
| *de novo* | 29,797 | 9168.03 | 1578.68 | 7.94 | 198.94 | 1094.31 |
| *Dreissena rostriformis* | 17,494 | 4,982.25 | 1,194.25 | 4.64 | 257.61 | 1,041.83 |
| *Sinonovacula constricta* | 15,305 | 6859.31 | 1,386.41 | 6.09 | 227.75 | 1,075.77 |
| mRNA | 102,143 | 5,095.79 | 1,103.53 | 2.99 | 369.01 | 2,005.64 |
| EVM | 25,174 | 9,103.73 | 1,534.29 | 7.47 | 205.48 | 1,170.51 |
| **Final set** | **25,135** | **9,112.83** | **1,535.27** | **7.47** | **205.41** | **1,170.40** |

**Table S4.** Gene set completeness measured by Benchmarking Universal Single-Copy Orthologs (BUSCO). Note: Busco was run in the “ -m protein” mode to access the gene set completeness, this is different form the genome completeness estimation in Table S1.

|  | **Number of**<br>**genes** | **Percent Completeness**<br>**(%)** |
| --- | --- | --- |
| Complete BUSCOs | 934 | 97.9 |
| Complete Single-copy<br>BUSCOs | 927 | 97.2 |
| Complete Duplicated<br>BUSCOs | 7 | 0.7 |
| Fragmented BUSCOs | 9 | 0.9 |
| Missing BUSCOs | 11 | 1.2 |

**Table S5.**
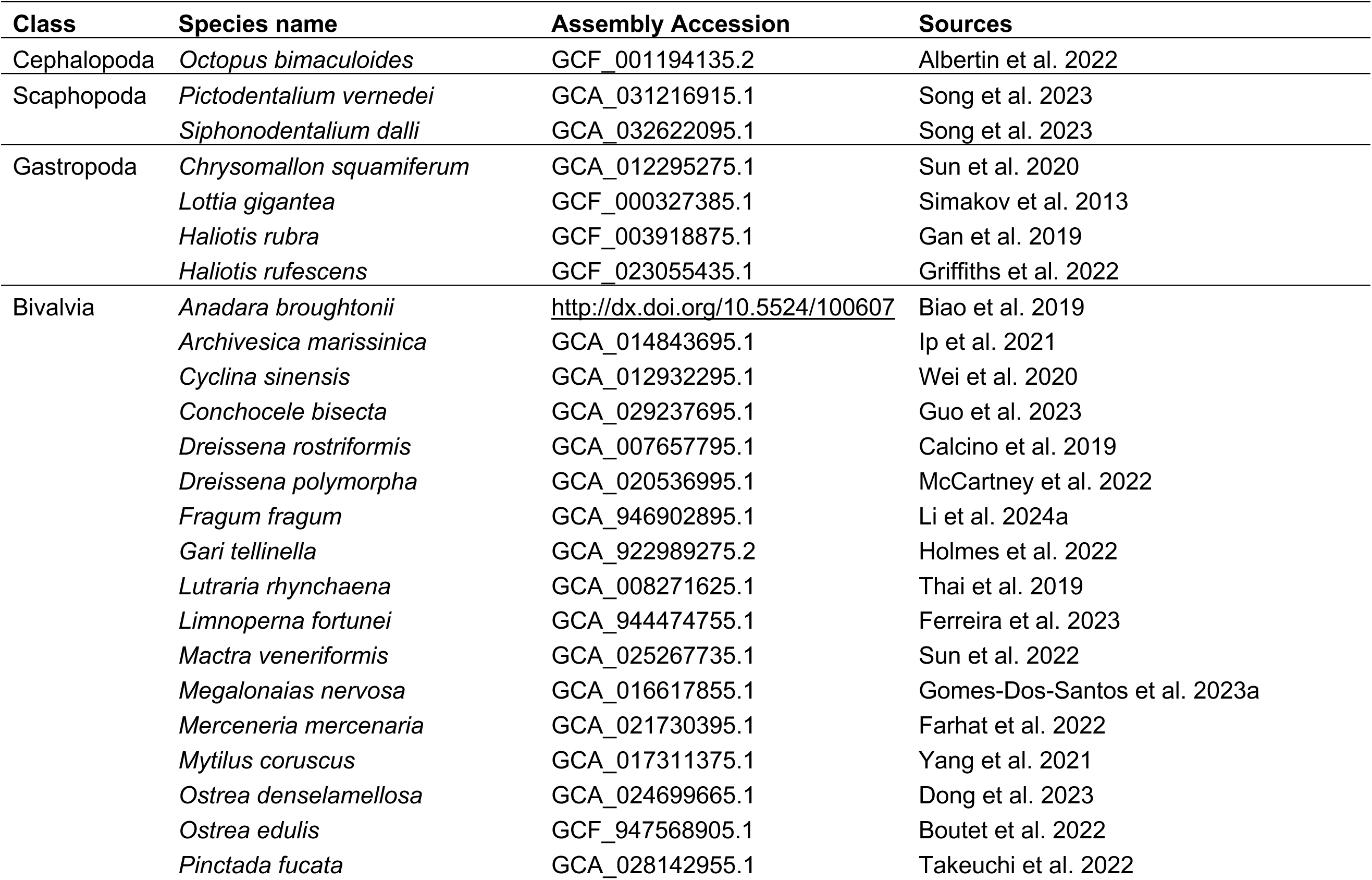
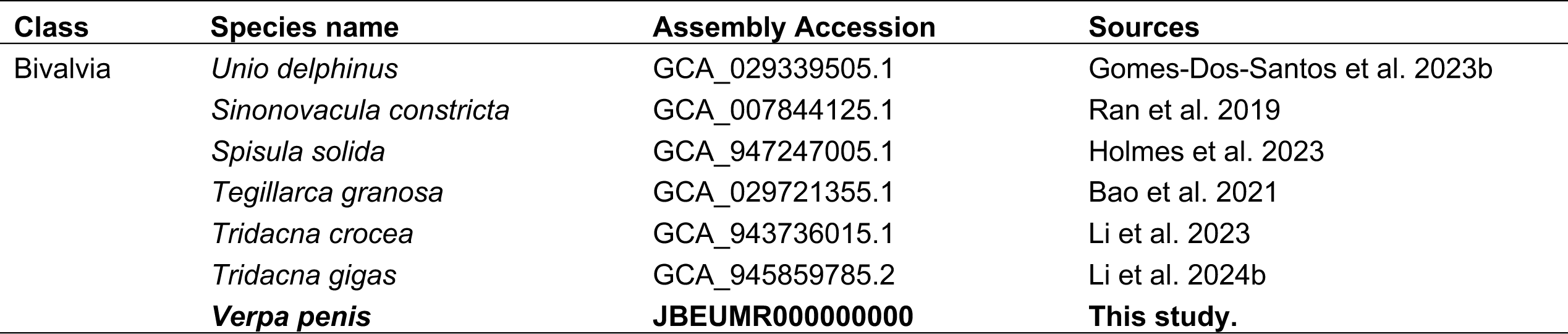
NCBI GenBank Accessions from publicly available data and newly sequenced *Verpa penis* (in bold) used for phylogenetic reconstruction.

## References

1. Ponder WF, Lindberg DR, Ponder JM. 2020. Biology and Evolution of the Mollusca. Boca Raton: CRC Press.

2. Sepkoski JJ. 1986 Phanerozoic Overview of Mass Extinction. In Patterns and Processes in the History of Life, pp. 277–295.

3. Collins KS, Edie SM, Jablonski D. 2023 Convergence and contingency in the evolution of a specialized mode of life: multiple origins and high disparity of rock-boring bivalves. Proc. R. Soc. B. 290, 20221907. (doi:10.1098/rspb.2022.1907)

4. Morton B, Machado FM. 2021 The origins, relationships, evolution and conservation of the weirdest marine bivalves: The watering pot shells. A review. In Advances in Marine Biology, pp. 137–220. Elsevier. (doi:10.1016/bs.amb.2021.03.001)

5. Morton, Brian. 2013 A cadaver unearthed: the anatomy of the Japanese living fossil *Stirpulina ramosa* (Bivalvia: Anomalodesmata: Clavagellidae) – the unique specimen in the collections of Emperor Shōwa. Zoological Journal of the Linnean Society (doi:10.1111/zoj12080)

6. Morton B. 2005 Biology and functional morphology of a new species of endolithic *Bryopa* (Bivalvia: Anomalodesmata: Clavagelloidea) from Japan and a comparison with fossil species of *Stirpulina* and other Clavagellidae. Invertebrate Biology 124, 202–219. (doi:10.1111/j.1744-7410.2005.00020.x)

7. Morton B. 2007 The evolution of the watering pot shells (Bivalvia: Anomalodesmata: Clavagellidae and Penicillidae). Rec West Aust Mus 24, 19. (doi:10.18195/issn.0312-3162.24(1).2007.019-064)

8. Machado FM, Passos FD. 2022 Revisiting the morphological aspects of the Anomalodesmata (Mollusca: Bivalvia): a phylogenetic approach. Invertebr. Syst. 36, 1063–1098. (doi:10.1071/IS22028)

9. Harper EM, Dreyer H, Steiner G. 2006 Reconstructing the Anomalodesmata (Mollusca: Bivalvia): morphology and molecules. Zoological Journal of the Linnean Society 148, 395–420. (doi:10.1111/j.1096-3642.2006.00260.x)

10. Williams ST, Foster PG, Hughes C, Harper EM, Taylor JD, Littlewood DTJ, Dyal P, Hopkins KP, Briscoe AG. 2017 Curious bivalves: Systematic utility and unusual properties of anomalodesmatan mitochondrial genomes. Molecular Phylogenetics and Evolution 110, 60–72. (doi:10.1016/j.ympev.2017.03.004)

11. Combosch DJ et al. 2017 A family-level Tree of Life for bivalves based on a Sanger-sequencing approach. Molecular Phylogenetics and Evolution 107, 191–208. (doi:10.1016/j.ympev.2016.11.003)

12. Dreyer H, Steiner G, Harper EM. 2003 Molecular phylogeny of Anomalodesmata (Mollusca: Bivalvia) inferred from 18S rRNA sequences. Zoological Journal of the Linnean Society 139, 229–246. (doi:10.1046/j.1096-3642.2003.00065.x)

13. Mayjonade B, Gouzy J, Donnadieu C, Pouilly N, Marande W, Callot C, Langlade N, Muños S. 2016 Extraction of High-Molecular-Weight Genomic DNA for Long-Read Sequencing of Single Molecules. BioTechniques 61, 203–205. (doi:10.2144/000114460)

14. Danecek P et al. 2021 Twelve years of SAMtools and BCFtools. GigaScience 10, giab008. (doi:10.1093/gigascience/giab008)

15. Baid G et al. 2022 DeepConsensus improves the accuracy of sequences with a gap-aware sequence transformer. Nat Biotechnol (doi:10.1038/s41587-022-01435-7)

16. Marçais G, Kingsford C. 2011 A fast, lock-free approach for efficient parallel counting of occurrences of *k* -mers. Bioinformatics 27, 764–770. (doi:10.1093/bioinformatics/btr011)

17. Ranallo-Benavidez TR, Jaron KS, Schatz MC. 2020 GenomeScope 2.0 and Smudgeplot for reference-free profiling of polyploid genomes. Nat Commun 11, 1432. (doi:10.1038/s41467-020-14998-3)

18. Cheng H, Concepcion GT, Feng X, Zhang H, Li H. 2021 Haplotype-resolved de novo assembly using phased assembly graphs with hifiasm. Nat Methods 18, 170–175. (doi:10.1038/s41592-020-01056-5)

19. Guan D, McCarthy SA, Wood J, Howe K, Wang Y, Durbin R. 2020 Identifying and removing haplotypic duplication in primary genome assemblies. Bioinformatics 36, 2896–2898. (doi:10.1093/bioinformatics/btaa025)

20. Li H. 2013 Aligning sequence reads, clone sequences and assembly contigs with BWA-MEM. arXiv:1303.3997

21. 2019 Picard Toolkit. http://broadinstitute.github.io/picard

22. Zhou C, McCarthy SA, Durbin R. 2023 YaHS: yet another Hi-C scaffolding tool. Bioinformatics 39, btac808. (doi:10.1093/bioinformatics/btac808)

23. Dudchenko O et al. 2018 The Juicebox Assembly Tools module facilitates *de novo* assembly of mammalian genomes with chromosome-length scaffolds for under $1000. (doi:10.1101/254797)

24. Durand NC, Robinson JT, Shamim MS, Machol I, Mesirov JP, Lander ES, Aiden EL. 2016 Juicebox provides a visualization system for Hi-C contact maps with unlimited zoom. Cell Systems 3, 99–101. (doi:10.1016/j.cels.2015.07.012)

25. Simão FA, Waterhouse RM, Ioannidis P, Kriventseva EV, Zdobnov EM. 2015 BUSCO: assessing genome assembly and annotation completeness with single-copy orthologs. Bioinformatics 31, 3210–3212. (doi:10.1093/bioinformatics/btv351)

26. Rhie A et al. 2021 Towards complete and error-free genome assemblies of all vertebrate species. Nature 592, 737–746. (doi:10.1038/s41586-021-03451-0)

27. Benson G. 1999 Tandem repeats finder: a program to analyze DNA sequences. Nucleic Acids Research 27, 573–580. (doi:10.1093/nar/27.2.573)

28. Tarailo-Graovac M, Chen N. 2009 Using RepeatMasker to Identify Repetitive Elements in Genomic Sequences. CP in Bioinformatics 25. (doi:10.1002/0471250953.bi0410s25)

29. Dobin A, Davis CA, Schlesinger F, Drenkow J, Zaleski C, Jha S, Batut P, Chaisson M, Gingeras TR. 2013 STAR: ultrafast universal RNA-seq aligner. Bioinformatics 29, 15–21. (doi:10.1093/bioinformatics/bts635)

30. Gabriel L, Brůna T, Hoff KJ, Ebel M, Lomsadze A, Borodovsky M, Stanke M. 2023 BRAKER3: Fully automated genome annotation using RNA-seq and protein evidence with GeneMark-ETP, AUGUSTUS and TSEBRA. (doi:10.1101/2023.06.10.544449)

31. Calcino AD, De Oliveira AL, Simakov O, Schwaha T, Zieger E, Wollesen T, Wanninger A. 2019 The quagga mussel genome and the evolution of freshwater tolerance. DNA Research 26, 411–422. (doi:10.1093/dnares/dsz019)

32. Dong Y et al. 2020 The chromosome-level genome assembly and comprehensive transcriptomes of the Razor Clam (*Sinonovacula constricta*). Front. Genet. 11, 664. (doi:10.3389/fgene.2020.00664)

33. Hauser M, Steinegger M, Söding J. 2016 MMseqs software suite for fast and deep clustering and searching of large protein sequence sets. Bioinformatics 32, 1323–1330. (doi:10.1093/bioinformatics/btw006)

34. Keilwagen J, Hartung F, Grau J. 2019 GeMoMa: Homology-Based Gene Prediction Utilizing Intron Position Conservation and RNA-seq Data. In Gene Prediction (ed M Kollmar), pp. 161–177. New York, NY: Springer New York. (doi:10.1007/978-1-4939-9173-0_9)

35. Grabherr MG et al. 2011 Full-length transcriptome assembly from RNA-Seq data without a reference genome. Nat Biotechnol 29, 644–652. (doi:10.1038/nbt.1883)

36. Haas BJ, Salzberg SL, Zhu W, Pertea M, Allen JE, Orvis J, White O, Buell CR, Wortman JR. 2008 Automated eukaryotic gene structure annotation using EVidenceModeler and the Program to Assemble Spliced Alignments. Genome Biol 9, R7. (doi:10.1186/gb-2008-9-1-r7)

37. Caballero M, Wegrzyn J. 2019 gFACs: Gene Filtering, Analysis, and Conversion to Unify Genome Annotations Across Alignment and Gene Prediction Frameworks. Genomics, Proteomics & Bioinformatics 17, 305–310. (doi:10.1016/j.gpb.2019.04.002)

38. Altenhoff AM et al. 2019 OMA standalone: orthology inference among public and custom genomes and transcriptomes. Genome Res. 29, 1152–1163. (doi:10.1101/gr.243212.118)

39. Katoh K, Standley DM. 2013 MAFFT Multiple Sequence Alignment Software Version 7: Improvements in Performance and Usability. Molecular Biology and Evolution 30, 772– 780. (doi:10.1093/molbev/mst010)

40. Katoh K. 2005 MAFFT version 5: improvement in accuracy of multiple sequence alignment. Nucleic Acids Research 33, 511–518. (doi:10.1093/nar/gki198)

41. Castresana J. 2000 Selection of Conserved Blocks from Multiple Alignments for Their Use in Phylogenetic Analysis. Molecular Biology and Evolution 17, 540–552. (doi:10.1093/oxfordjournals.molbev.a026334)

42. Cunha TJ, Giribet G. 2019 A congruent topology for deep gastropod relationships. Proc. R. Soc. B. 286, 20182776. (doi:10.1098/rspb.2018.2776)

43. Smith SA, Dunn CW. 2008 Phyutility: a phyloinformatics tool for trees, alignments and molecular data. Bioinformatics 24, 715–716. (doi:10.1093/bioinformatics/btm619)

44. Minh BQ, Schmidt HA, Chernomor O, Schrempf D, Woodhams MD, Von Haeseler A, Lanfear R. 2020 IQ-TREE 2: New Models and Efficient Methods for Phylogenetic Inference in the Genomic Era. Molecular Biology and Evolution 37, 1530–1534. (doi:10.1093/molbev/msaa015)

45. Kalyaanamoorthy S, Minh BQ, Wong TKF, Von Haeseler A, Jermiin LS. 2017 ModelFinder: fast model selection for accurate phylogenetic estimates. Nat Methods 14, 587–589. (doi:10.1038/nmeth.4285)

46. Hoang DT, Chernomor O, Von Haeseler A, Minh BQ, Vinh LS. 2018 UFBoot2: Improving the Ultrafast Bootstrap Approximation. Molecular Biology and Evolution 35, 518–522. (doi:10.1093/molbev/msx281)

47. WoRMS Editorial Board. 2024 World Register of Marine Species. Available from https://www.marinespecies.org at VLIZ. Accessed yyyy-mm-dd. (doi:10.14284/170)

48. Morton B, Machado FM. 2019 Predatory marine bivalves: A review. 84, 1–98.

49. Gonzalo Giribet, Distel, D.L. 2003 Bivalve phylogeny and molecular data. In Molecular Systematics and Phylogeography of Mollusks (eds Lydeard, C., Lindberg, David R.), pp. 45–90. Washington, D.C.: Smithsonian Institution, Washington, D.C.

50. Sharma PP et al. 2012 Phylogenetic analysis of four nuclear protein-encoding genes largely corroborates the traditional classification of Bivalvia (Mollusca). Molecular Phylogenetics and Evolution 65, 64–74. (doi:10.1016/j.ympev.2012.05.025)

51. Bieler R et al. 2014 Investigating the Bivalve Tree of Life – an exemplar-based approach combining molecular and novel morphological characters. Invert. Systematics 28, 32. (doi:10.1071/IS13010)

52. Sigwart JD, Lindberg DR, Chen C, Sun J. 2021 Molluscan phylogenomics requires strategically selected genomes. Philosophical Transactions of the Royal Society B: Biological Sciences 376, 20200161. (doi:10.1098/rstb.2020.0161)

53. Sigwart JD, Li Y, Chen Z, Voncina K, Sun J. 2024 Still waters run deep: Large scale genome rearrangements in the evolution of morphologically conservative Polyplacophora. (doi:10.1101/2024.06.13.598811)

54. Jenkinson JJ. 2014 Chromosomal characteristics of North American and other naiades (Bivalvia: Unionida). Malacologia 57, 377–397. (doi:10.4002/040.057.0210)

55. Guo Y et al. 2023 Hologenome analysis reveals independent evolution to chemosymbiosis by deep-sea bivalves. BMC Biol 21, 51. (doi:10.1186/s12915-023-01551-z)

56. Li R et al. 2024 The genome sequence of the giant clam, *Tridacna gigas* (Linnaeus, 1758). Wellcome Open Res 9, 145. (doi:10.12688/wellcomeopenres.21136.1)

57. Li R et al. 2024 The genome sequence of a heart cockle, *Fragum fragum* (Linnaeus, 1758). Wellcome Open Res 9, 129. (doi:10.12688/wellcomeopenres.21134.1)

58. Nosil P, Soria-Carrasco V, Villoutreix R, De-la-Mora M, De Carvalho CF, Parchman T, Feder JL, Gompert Z. 2023 Complex evolutionary processes maintain an ancient chromosomal inversion. Proc. Natl. Acad. Sci. U.S.A. 120, e2300673120. (doi:10.1073/pnas.2300673120)

59. Liu Z, Roesti M, Marques D, Hiltbrunner M, Saladin V, Peichel CL. 2022 Chromosomal fusions facilitate adaptation to divergent environments in Threespine Stickleback. Molecular Biology and Evolution 39, msab358. (doi:10.1093/molbev/msab358)

60. Wilson J, Staley JM, Wyckoff GJ. 2020 Extinction of chromosomes due to specialization is a universal occurrence. Sci Rep 10, 2170. (doi:10.1038/s41598-020-58997-2)

61. Savazzi E. 2000 Morphodynamics of *Bryopa* and the evolution of clavagellids. SP 177, 313–327. (doi:10.1144/GSL.SP.2000.177.01.20)

62. Purchon RD. 1956 A note on the biology of *Brechites penis* (L.) (Lamellibranchia). Journal of the Linnean Society of London, Zoology 43, 43–54. (doi:10.1111/j.1096-3642.1956.tb02506.x)

63. Purchon, R.D. In press. A further note on the biology of Brechites penis (L.) (Lamellibranchia). Proceedings of the Malacological Society of London 34, 19–23.

64. Savazzi E. 1982 Adaptations to tube dwelling in the Bivalvia. Lethaia 15, 275–297. (doi:10.1111/j.1502-3931.1982.tb00650.x)

65. Vermeij GJ, Thomson TJ. 2023 Imbricated shell sculpture in benthic bivalves. Journal of Morphology 284, e21564. (doi:10.1002/jmor.21564)

66. Vermeij GJ. 2022 The balanced life: evolution of ventral shell weighting in gastropods. Zoological Journal of the Linnean Society 194, 256–275. (doi:10.1093/zoolinnean/zlab019)

67. Morton B. 2002 The biology and functional morphology of *Humphreyia strangei* (Bivalvia: Anomalodesmata: Clavagellidae): an Australian cemented ‘watering pot’ shell. Journal of Zoology 258, 11–25. (doi:10.1017/S0952836902001164)

68. Tan SK, Tan SH, Low MEY. 2011 A reassessment of *Verpa penis* (Linnaeus, 1758) (Mollusca: Bivalvia: Clavagelloidea), a species presumed nationally extinct. Nature in Singapore. 4, 5–8.

